# Genome Sequence of Indian Peacock Reveals the Peculiar Case of a Glittering Bird

**DOI:** 10.1101/315457

**Authors:** Shubham K. Jaiswal, Ankit Gupta, Rituja Saxena, P. K. Vishnu Prasoodanan, Ashok K. Sharma, Parul Mittal, Ankita Roy, Aaron B.A. Shafer, Nagarjun Vijay, Vineet K. Sharma

**Affiliations:** Metagenomics and Systems Biology Group, Department of Biological Sciences, Indian Institute of Science Education and Research Bhopal, India; Computational Evolutionary Genomics Lab, Department of Biological Sciences, Indian Institute of Science Education and Research Bhopal, India; Forensic Science and Environmental & Life Sciences, Trent University, Canada

**Keywords:** Peacock genome, Peafowl, Comparative genomics, dN/dS, positive selection, Adaptive evolution, Hamilton-Zuk hypothesis

## Abstract

The unique ornamental features and extreme sexual traits of Peacock have always intrigued the scientists. However, the genomic evidence to explain its phenotype are yet unknown. Thus, we report the first genome sequence and comparative analysis of peacock with the available high-quality genomes of chicken, turkey, duck, flycatcher and zebra finch. The candidate genes involved in early developmental pathways including TGF-β, BMP, and Wnt signaling pathway, which are also involved in feather patterning, bone morphogenesis, and skeletal muscle development, showed signs of adaptive evolution and provided useful clues on the phenotype of peacock. The innate and adaptive immune components such as complement system and T-cell response also showed signs of adaptive evolution in peacock suggesting their possible role in building a robust immune system which is consistent with the between species predictions of Hamilton-Zuk hypothesis. This study provides novel genomic and evolutionary insights into the molecular understanding towards the phenotypic evolution of Indian peacock.

## INTRODUCTION

One of the most glittering bird, the Indian peafowl (*Pavo cristatus*), is an avian species that had once puzzled the greatest naturalist, Charles Darwin, who wrote - “the sight of a feather in a Peacock’s tail, whenever I gaze at it, makes me sick” (Huxley, 1968). The presence of an exceptional ornamental plumage with large tail-coverts in peacock, which makes it more visible to predators attack, posed a question for his theory of natural selection. However, later studies showed its significance for the reproductive success of peacock mediated by sexual selection. The *Pavo* genus from the family Phasianidae has two known species, *Pavo cristatus* (Blue peafowl) and *Pavo muticus* (Green peafowl), which diverged about 3 million years ago (OUYANG et al., 2009). The Blue peafowl (Indian Peacock) is endemic to the Indian subcontinent, whereas, the Green Peafowl is mostly found in Southeast Asia.

Peacock (male peafowl) is one of the largest known bird among pheasants and flying birds. It shows sexual dimorphism, polygamy with no paternal care to offspring, and an elaborate male display for mating success (Zahavi, 1975;Ramesh and McGowan, 2009). The sexual selection is extreme in peacock, which is dependent upon the ornamental display (glittering train and crest plumage) and behavioral traits (Loyau et al., 2005a). These ornamental features are also used as an honest signal about their immunocompetence to the peahen, which helps in the selection of individuals with better immunity (Loyau et al., 2005b). Though, the male masculine traits are testosterone-dependent in peacock, the large train is the default state since the peahen also shows the development of this train after ovirectomy(Owens and Short, 1995).

The existence of intricate ornaments in peacock has perplexed the scientists for decades and has led to several ecological and population-based studies (Zahavi, 1975;Loyau et al., 2005a;Ramesh and McGowan, 2009). However, the genomic details about the phenotypic evolution of this species are still unknown. Therefore, we carried out the comprehensive comparative genomics of *Pavo cristatus* (Blue Peafowl) to decipher the genomic evolution of this species. The ornamental and sexual characteristics of peacock are distinct from other birds and are absent in the available closely related species such as chicken and turkey, which makes it intriguing to look for the genomic changes underlying the phenotypic divergence of peacock. Therefore, we also carried out a comprehensive comparative genome-wide analysis of peacock genome (order Galliformes) with the high quality genomes of five other birds under the class Aves: chicken and turkey (order Galliformes), duck (order Anseriformes), and flycatcher and zebra finch (order Passeriformes). The comparative genome-wide analysis of peacock with five other related birds provided novel genomic insights into the intriguing peacock genome evolution.

## MATERIALS AND METHODS

### Sample collection, DNA isolation, and sequencing of peacock genome

Approximately 2 ml blood was drawn from the medial metatarsal vein of a two years old male Indian peacock at Van Vihar National Park, Bhopal, India and was collected in EDTA-coated vials. The fresh blood sample was immediately brought to the laboratory at 4 °C and genomic DNA was extracted using DNeasy Blood and Tissue Kit (Qiagen, USA) following the manufacturer’s protocol. Sex of the bird was determined to be male by morphological identification and was confirmed using molecular sexing assay (Supplementary Note). Multiple shotgun genomic libraries were prepared using Illumina TruSeq DNA PCR-free library preparation kit and Nextera XT sample preparation kit (Illumina Inc., USA), as per the manufacturer’s protocol. The insert size for the TruSeq libraries was selected to be 550 bp and the average insert size for Nextera XT libraries was ~650 bp. The sequencing library size for both the libraries was assessed on 2100 Bioanalyzer using High Sensitivity DNA kit (Agilent, USA). The libraries were quantified using KAPA SYBR FAST qPCR Master mix with Illumina standards and primer premix (KAPA Biosystems, USA), and Qubit dsDNA HS kit on a Qubit 2.0 fluorometer (Life Technologies, USA) as per the Illumina suggested protocol. The normalised libraries were loaded on Illumina NextSeq 500 platform using NextSeq 500/550 v2 sequencing reagent kit (Illumina Inc., USA) and 150 bp paired-end sequencing was performed for all the libraries on May 11, 2016.

#### Sequence alignment and phylogenetic tree construction

All sequence alignments (DNA and Protein) used for the phylogenetic tree reconstruction and other sequence divergence analysis were generated using MUSCLE release 3.8.31 (Edgar, 2004). The likelihood-based tree-searching algorithm was used for phylogenetic tree reconstruction using PhyML version 3.1 (Guindon et al., 2010). For nucleotide sequences GTR model was used, whereas for protein sequences JTT model was utilized. The bootstrapping value of n=1000 was used to test the robustness of the constructed phylogenetic trees of mitochondrial genome and concatenated nuclear-genes.

#### Gene gain/loss analysis

To estimate the gene gain and loss in gene families, CAFE (v3.1)(Han et al., 2013) with a random birth and death model was used (Supplementary Figure 7). The species tree was constructed using NCBI taxonomy, and the branch lengths were calculated using the fossil data in TimeTree as described in Ensembl Compara pipeline(Vilella et al., 2009). Simulated data based on the properties of observed data was generated using the command gene family and the significance of two-lambda model (separate lambda values for Galloanserae) was assessed against a global lambda model. The two-parameter model was found to fit the data better because the observed LR [LR = 2*(score of global lambda model – score of multi-lambda model)] was greater than 95% of the distribution of simulated LRs. Therefore, the two-lambda model was used for the following CAFE analysis.

#### Identification of CDS with multiple signs of adaptive evolution

All validated peacock coding gene sequences (>90% valid bases) were analyzed through multiple sequence-based analysis such as dN/dS or ω (ratio of the rate of non-synonymous to the rate of synonymous substitutions) enrichment, positive selection, and unique substitution to assess the adaptive sequence divergence. The functional analysis was performed using KEGG (Kanehisa and Goto, 2000), eggNOGs (Huerta-Cepas et al., 2016) and NCBI NR (O’Leary et al., 2016) databases. Furthermore, the functional impact of the identified unique substitutions and other sequence variations were evaluated using functional domain analysis and SIFT (Sorting Intolerant from tolerant) analysis (Kumar et al., 2009). SIFT is a homology-based method, where the specific-amino acids of protein sequence conserved across species are considered to be functionally crucial.

##### dN/dS enrichment analysis

Based on the dN/dS or ω values, the positively selected (ω >1), negatively selected (ω <1) and neutrally selected (ω =1) genes were identified. The dN/dS values for the peacock CDS were calculated using CODEML program of the PAML package 4.9(Yang, 2007). The pairwise dN/dS analysis was performed on the orthologous genes for six different pairs: peacock-chicken, peacock-turkey, peacock-duck, peacock-zebra finch, peacock-flycatcher and chicken-turkey using default parameters. To check for the convergence of calculated values, the iterations were performed with three different initial or fixed ω values, i.e. 0.5, 1 and 1.5, and only the coding gene sequences with consensus values were considered. To reduce the false positives and aberrant dN/dS values from analysis all the genes with the dN/dS values above five were not used for the function interpretation of results and for drawing conclusions out of it, although, they were used at the eggNOG and KEGG functional classification stage to reduce the bias.

##### Positive selection analysis

The multiple sequence alignment for each peacock coding gene sequence and the corresponding orthologs identified using reciprocal blast approach in the other five bird genomes were carried out using EMBOSS tranalign program (Rice et al., 2000). Furthermore, the Maximum Likelihood-based (ML) phylogenetic tree was constructed using the amino acid sequence of these orthologs. Based on the alignment and the phylogenetic tree, the calculations of likelihood scores with revised branch-site model A was performed to identify the signatures of positive selection in peacock for the considered coding gene sequence. This model tries to detect positive selection acting on specific sites on the particular specified branch (foreground branches) (Yang et al., 2005;Zhang et al., 2005). The foreground branch consisted of peacock, and the other branches constituted the ‘background branches’. Codons were categorized into previously assumed four classes in the model based on the foreground and background estimates of dN/dS (ω) values. The alternative hypothesis, according to which the foreground branches show positive selection with ω >1, was compared with the null hypothesis, according to which all branches have the same ω =1 value. The comparison was performed using LRT (Likelihood Ratio Test) values based chi-square test. The genes with P-value < 0.05 were considered to be positively selected in peacock. Additionally, the amino acid sites under positive selection were identified using the Bayesian Empirical Bayes values for the branch-site model A (Zhang et al., 2005). This positive selection analysis was performed using CODEML program of the PAML package version 4.9 (Yang, 2007).

##### Unique substitution analysis

The peacock coding gene sequence and its orthologs identified from the five bird genomes were translated using EMBOSS transeq and the protein sequence alignments were performed using MUSCLE release 3.8.31 (Edgar, 2004). Using custom-made Perl scripts, the positions at which the peacock protein showed amino acid substitutions in comparison to all the other five bird genomes were identified and reported as the unique substitutions in peacock genome.

#### Estimation of effective population size (N_e_) history

The demographic history of the peacock was reconstructed by estimating the effective population size (N_e_) over time using pairwise sequentially Markovian coalescent (PSMC) (Li and Durbin, 2011). The autosomal data of the peacock diploid genome sequence was filtered by excluding sites at which the inferred consensus quality was below 20, and the read depth was either one-third or more than twice of the average read depth across the genome. Since, mean coverage and percentage of missing data, both are important filtering thresholds in PSMC analysis, the minimum length of the contigs selected for carrying out the analysis was 5000 bp based on no more than 25% of the missing data as suggested by Krystyna et al. (Nadachowska-Brzyska et al., 2016) (**Supplementary Figure 5**). The resultant filtered genome sequence used for the analysis was 76% of the total genome. The parameters for PSMC were set to “N30 -t5 -r5 -p4+30*24+610”, which were used previously for 38 bird species (Nadachowska-Brzyska et al., 2015). Generation time and mutation rate are necessary to scale the results of PSMC analysis to real time. Hence, a generation time of 4 years was used in this analysis and was calculated as twice of the sexual maturity (2 years) [26]. The *mutation rate of 1.33e-09 was used as calculated in a previous study (Wright et al., 2015). It is known that the estimates of Ne from PSMC can be influenced by the quality of the genome and sequencing coverage. To ensure that our results are not strongly influenced by such artefacts, 100 bootstrap runs were performed to estimate the N_e_ from different parts of the genome to ascertain variability in the estimates of N_e_.

#### Accession codes

Sequence data for *Pavo cristatus* has been deposited in Short Read Archive under project number SRP083005 (BioProject accession: PRJNA040135, Biosample accession: SAMN05660020) and accession codes : SRR4068853 and SRR4068854.

## RESULTS

Although more than fifty bird genomes have been sequenced so far, yet the comprehensive and curated gene set is available only for the handful of bird genomes at the Ensembl browser. Thus, the comparative genomics analysis was performed using only the high quality genome assemblies of species relatively closer to pheasants which were available at the Ensembl browser.

The whole genome sequencing of peacock genome yielded 153.7 Gb of sequence data (~136x genomic coverage; **Supplementary Table 1 and Supplementary Figure 1 and 2**). High-quality sequence reads were used to generate a draft genome assembly of an estimated genome size of 1.13 Gb using Abyss, Gapcloser, and Agouti (**Supplementary Table 2**). The de novo genome scaffold and contig N50s were 25.6 Kb and 19.3 Kb, respectively (**Supplementary Table 2**). BUSCO scores assessed the genome assembly to be 77.6% complete (S:63.44%, D:14.2%) and predicted 13.5% as partial, and 8.9% as missing BUSCOs (**Supplementary Table 3**). Using ab initio-based approach, 25,963 coding sequences were identified in peacock, and in addition, 213 tRNAs, 236 snoRNAs, and 540 miRNAs were also identified (**Supplementary Table 4**). The peacock genome was found to have less repetitive DNA (8.62%) as compared to chicken (9.45%) (**Supplementary Table 5**). PSMC analysis suggested that the peacock suffered at least two bottlenecks (around four Million and 450,000 years ago), which resulted in a severe reduction in its effective population size (**Figure 1**). It was also interesting to note that the results of PSMC analysis of peacock were similar to the demographic history of the tropical bowerbird and turkey vulture that show long-term decrease in the effective population size (Nadachowska-Brzyska et al., 2015), perhaps because all three birds are native to the tropical rain forests.

**Figure 1:**
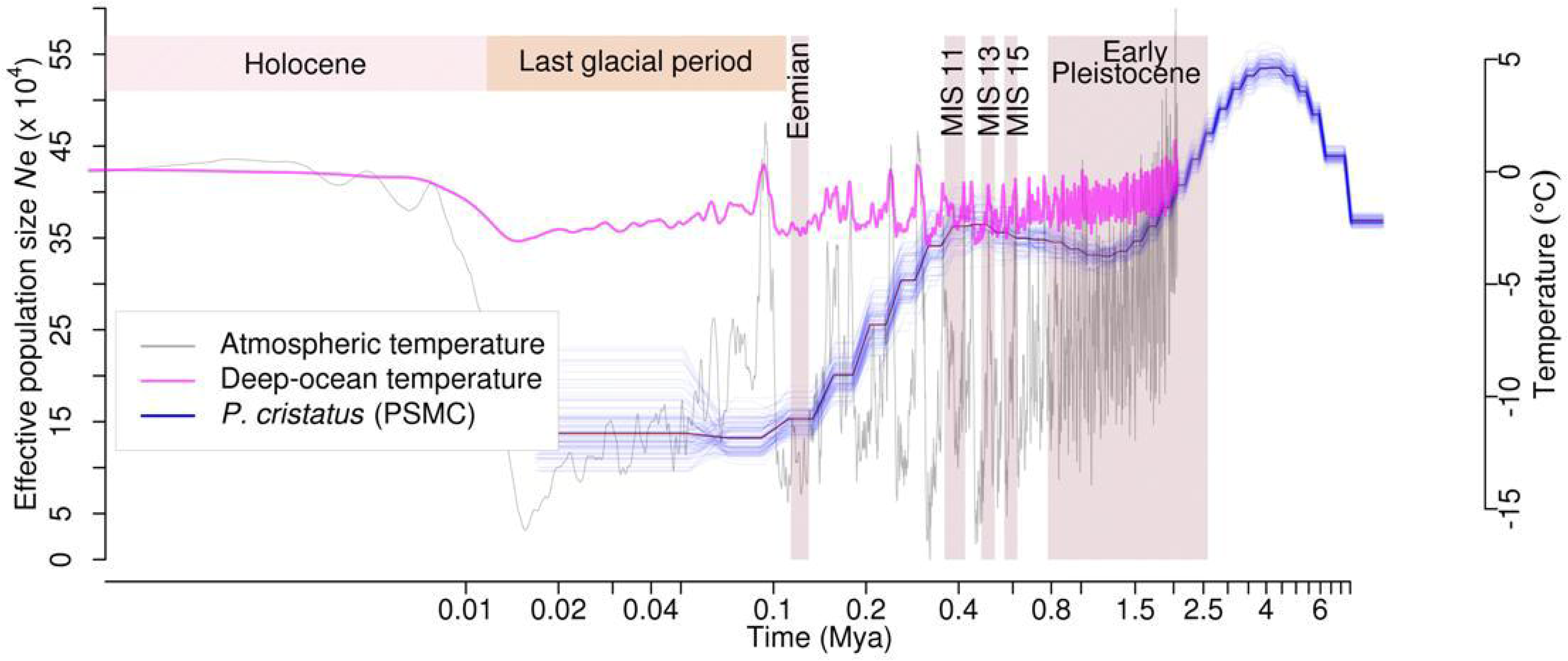
Effective population size (N_e_) estimated from PSMC analysis for Peacock. The changes in effective population size (N_e_) for the peacock is shown as the blue line plot. The thick line represents the consensus, and the thin light line corresponds to 100 bootstrapping rounds. Atmospheric and deep ocean temperatures from (Bintanja and Van de Wal, 2008) have been overlaid.

Using a combination of homology and *ab initio* based approaches, 15,970 protein-coding genes were identified in peacock by utilizing the peacock genome assembly and the filtered high quality reads from previous study (**Supplementary Methods**). The comparison of single nucleotide variants (SNVs) between chicken and peacock revealed 2,051,161 heterozygous SNVs at a rate of 2.05 SNV per Kb. The observed SNV rate in peacock was closer to turkey in comparison to the other avian species (**Supplementary Note and Supplementary Table 6**).

The analysis of gene gain/loss in gene families was also performed for the six bird genomes namely peacock, chicken, turkey, duck, flycatcher and zebra finch. The Venn diagram of the genes families for these bird genomes is shown in **Figure 2A**. Additionally, the phylogenetic tree showing the gene gain/loss for the six bird genomes and the outlier green anole is displayed in **Figure 2B**. It is apparent that the common ancestor to the birds in the phylogenetic tree show a loss of 2,295 genes, which is also supported by a previous report mentioning the loss of around 2000 genes in the ancestor as compared to other vertebrate lineages (Huang et al., 2013;Lovell et al., 2014). However, such observations could be an artefact of poor genome coverage in the GC-rich regions and incomplete genome assemblies (Bornelöv et al., 2017). This can also lead to an over or under-estimation of gene counts due to fragmentation of genes on multiple contigs and gaps in the assembly (Denton et al., 2014). We observed that contraction has been more prominent in comparison to expansion for the common ancestor of Galliformes and Anseriformes and the same pattern has also been observed for turkey and duck (**Figure 2B**). These observations corroborates with the previous study (Huang et al., 2013). However, an opposite pattern of expansion in gene families was observed for peacock and chicken (**Figure 2B**). The top 20 protein families featuring gain and loss in the peacock genome are listed in **Supplementary Table 7, 8 and 9**.

**Figure 2:**
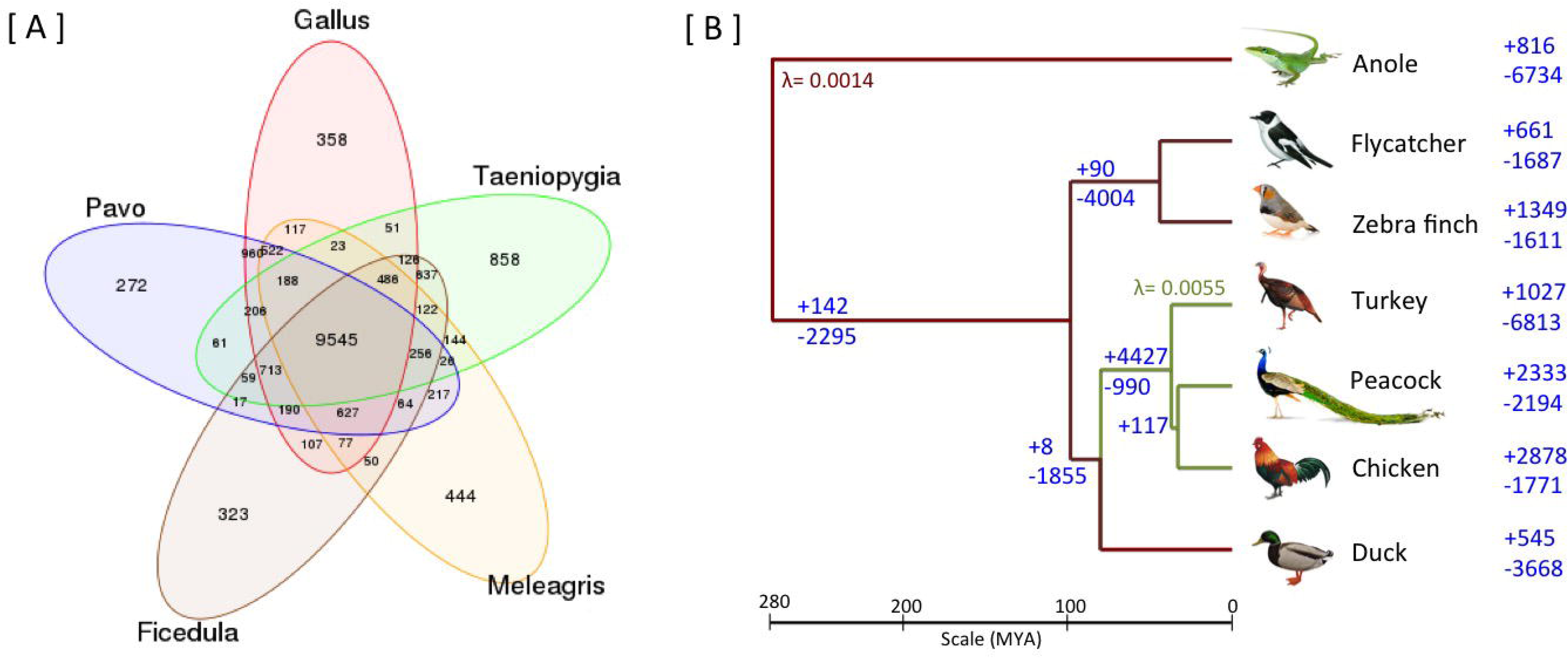
**[A] Venn diagram of gene families identified using TreeFam.** A total of 9,545 gene families were common among the five bird genomes. 522 gene families were unique to the genus (Pavo, Gallus and Meleagris) of Galliformes order, whereas, 637 gene families were unique to the genus (Ficedula and Taeniopygia) of Passeriformes order. **[B] Gene gain/loss in the six avian species and anole** The number of gene gain (+) and loss (−) are mentioned on the right of the taxa (branches), for the six avian species and an outlier green anole. The gene gain and loss were calculated using CAFE two-lambda model with λ = 0.0055 for Galliformes and λ= 0.0014 for the rest of the tree.

The phylogenetic position of peacock was determined using a maximum likelihood-based analysis performed using the coding sequences of 5,907 orthologous genes identified from the six bird genomes : peacock, chicken, turkey, duck, flycatcher and zebra finch genomes (**Supplementary Note**). From the phylogenetic tree, it was apparent that peacock is closer to chicken than turkey in the Galliformes order, and formed a monophyletic group with duck from Anseriformes order (**Figure 3A).** The genome-wide analysis confirms the earlier studies carried out using limited coding and non-coding sequences, and chromosomal banding patterns (Stock and Bunch, 1982;Kaiser et al., 2007;Wang et al., 2013). The branch-specific ω or dN/dS (ratio of the rate of non-synonymous to synonymous substitutions) values were lower for chicken and peacock in comparison to the other bird genomes (**Figure 3A**). The mitochondrial genome, which evolves independent of the nuclear genome, was also used to infer the phylogenetic relationships using the complete mitochondrial genome sequences of peacock and 22 species from five different classes of Chordates, which included Aves, Mammalia, Reptilia, Actinopterygii and Amphibia (**Supplementary Figure 3**). The phylogenetic positions of the six bird species were found similar in both the trees (**Figure 1A, Supplementary Figure 4**). Furthermore, the distribution of ω values and log-transformed mean ω values for the 5,907 orthologous genes showed the evolutionary closeness of peacock and chicken in comparison to peacock and turkey and supported the observations made from the phylogenetic trees (**Figure 3B**). The phylogenetic analysis carried out using nuclear-genes and mitochondrial genomes revealed that peacock is closer to chicken as compared to turkey, which confirms the phylogenetic position of peacock through a genome-wide analysis, in addition to the earlier reports from limited molecular data (Stock and Bunch, 1982;Kimball et al., 1999;Kan et al., 2010;Wang et al., 2013).

**Figure 3:**
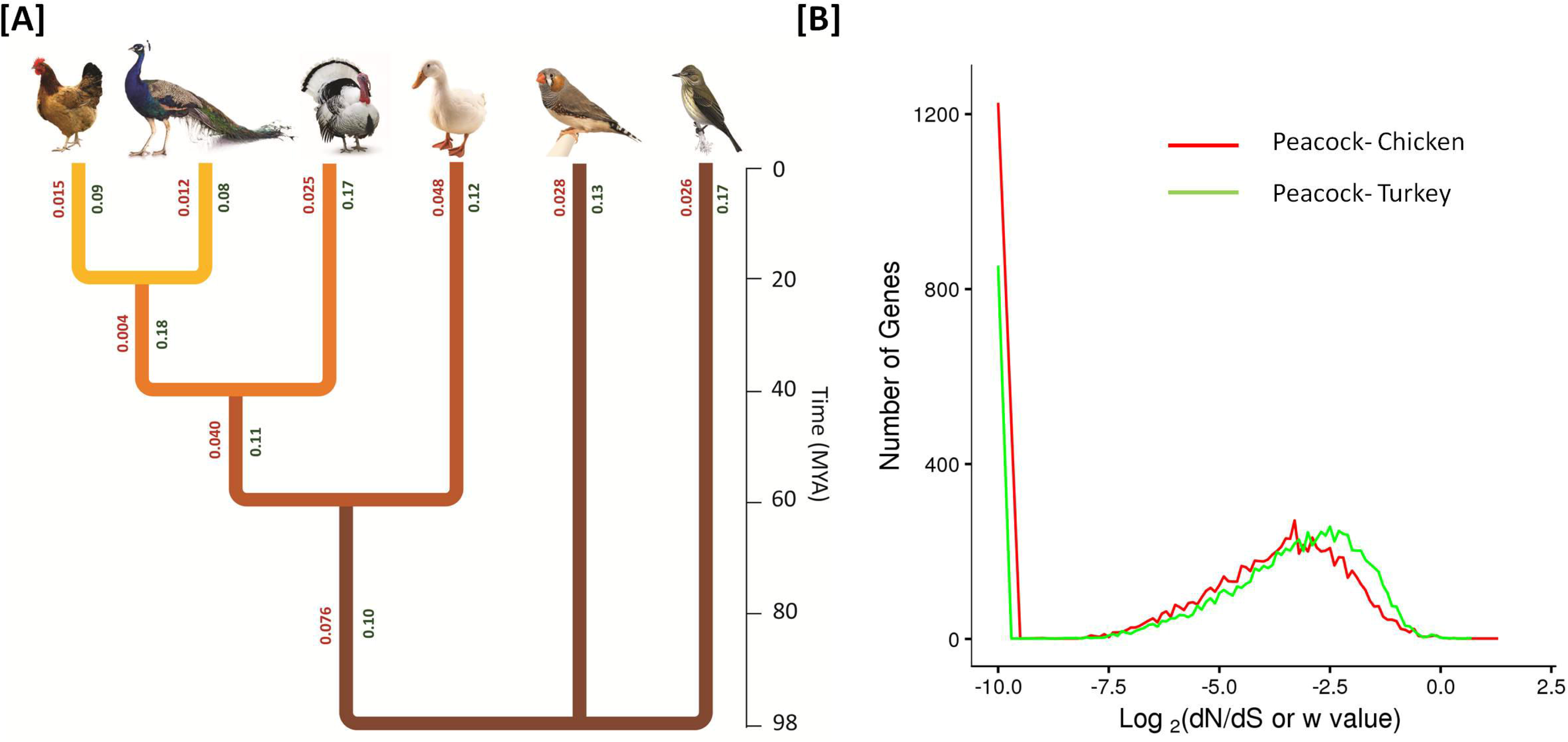
**[A] Phylogenetic relationship of peacock with other bird genomes** The phylogenetic tree constructed from the concatenated alignments of the orthologous genes across all six species. The divergence time of different bird species was determined using the TIMETREE database (Hedges et al., 2006), which is based on the published reports of molecular and fossil data. The origin of turkey was estimated to be 37.2 mya, whereas the origin of peacock and chicken was estimated to be 32.9 mya. **[B] Comparison of the distribution of ω or dN/dS values for the pairs of birds in Galliformes order: peacock-chicken (PG) and peacock-turkey (PT).** The calculation was performed using 9,078 orthologous genes by employing CODEML program of PAML package v4.9a. The actual values were log-transformed to the base of 2 and mean values for the PG and PT pairs were -4.4 and -3.8, respectively.

### Divergence and adaptive evolution

A comparative genomic analysis was performed using 15,970 peacock genes and their corresponding orthologs present in chicken, turkey, duck, flycatcher and zebra finch. The dN/dS values >1 was shown by 74 genes, of which 25 genes had values above five indicating possible false positives, and were not considered for the functional interpretation (**Supplementary Table 10**). A total of 491 genes displayed the signs of positive selection identified using branch-site model A and the statistical significance was evaluated using likelihood ratio tests with p-value threshold of 0.05 (**Supplementary Table 11**). Unique amino-acid substitutions in peacock were found for 3,238 genes, of which the substitutions in 116 genes were predicted to affect the protein function using SIFT (Sorting Intolerant from Tolerant) analysis (**Supplementary Table 12 and 13**). A total of 417 genes contained amino acid sites, which were under significant positive selection based on the Bayesian empirical Bayes values. In total 99 genes showed positive selection and unique amino acid substitutions that may affect the protein function predicted using SIFT and are referred to as genes with ‘multiple signs of adaptation’ (MSA) in this study (**Supplementary Table 14**).

The functional analysis revealed the role of these genes in key cellular processes such as cell proliferation and differentiation (MAPK, RAS, PI3K-Akt, ErbB, Hippo, Rap1, and Jak-STAT signaling, Wnt signaling, calcium signaling and adrenergic signaling in cardiomyocytes) and immune response (T cell receptor, Toll-like receptor signaling, NOD-like receptor signaling, complement and coagulation cascade and chemokine-chemokine signaling). In addition, multiple genes involved in early development pathways such as TGF-β, Wnt/β-catenin, FGF, and BMP signaling also showed adaptive sequence divergence in peacock. These cellular processes and pathways regulate key features such as early development, feather development, bone morphogenesis, skeletal muscle development, metabolism, and immune response (**Supplementary Table 15**).

An interesting observation was made from the signalling pathways such as Wnt, Rap1, Ras, Jak-Stat, and cAMP-mediated GPCR signalling. It was observed that the ligand and/or receptor, and in some cases the final effector genes showed adaptive evolution, whereas the genes involved in the intermediate signal transduction processes remained conserved perhaps due to their common role in multiple signaling pathways (**Supplementary Note**). Another interesting observation was that in several interacting protein pairs, both the interacting proteins showed sequence divergence hinting towards their co-evolution (Moyle et al., 1994). These protein pairs were majorly involved in early development pathways such as Wnt, BMP and TGF-β signalling, cell cycle regulation, DNA replication, GPCR signaling, and gene expression regulation (**Supplementary Table 16**).

### Adaptive evolution of early developmental pathways

The early developmental pathways, which are crucial in guiding the embryonic development in birds such as TGF-β, Wnt, FGF and BMP signaling, showed adaptive divergence in peacock (Klaus and Birchmeier, 2008). Among these pathways, the TGF-β pathway is known to regulate the cartilage connective tissue development (Loveridge et al., 1993), and also functions as an activator of feather development in birds. In this pathway, TGFBR3 gene showed MSA, and TGF-β3 preproprotein, TGFBRAP1, and TAB3 genes showed multiple unique substitutions (**Supplementary Note**). The Wnt signaling pathway is involved in development, regeneration, aging process (Brack et al., 2007;Klaus and Birchmeier, 2008), and also regulates the initial placement of feather buds and their consolidation within the feather field (Lim and Nusse, 2013). Multiple regulators of Wnt signaling such as WNT2, WIF1, and DKK2 genes had positively selected amino acid sites and showed signs of adaptive evolution. The WIF1 and DKK2 genes also harbored multiple unique substitutions. Furthermore, the DKK2 and WNT2 genes were found to be positively selected in peacock. APCDD1 gene, which is an inhibitor of Wnt signaling pathway, showed MSA. The Bone Morphogenetic Protein (BMP) signaling is involved in the development of skeletal muscles, bone and cartilage connective tissue (Nie et al., 2006;Nishimura et al., 2012), neurogenesis (Groppe et al., 2002), and feather formation and patterning. Multiple genes such as BRK-3, BMP5, BMP3, BMP10 and CRIM1, which are involved in the regulation of BMP pathways and the corresponding early development, showed unique substitutions that may affect their function in cellular pathways as compared to the other birds.

In addition, the Notch-2 receptor gene of Notch-Delta signaling, which is involved in growth and patterning of feather buds, early development of sensory organs (Crowe et al., 1998), and terminal muscle differentiation also showed five unique substitutions. Unique substitutions were also found in the FGFR3 receptor gene and FGF23 genes, which are part of the FGF signaling involved in limb and skeletal muscle development, feather development and morphogenesis, and regulation of feather density and patterning (Pownall and Isaacs, 2010).

Taken together, the multiple signs of evolution observed in the genes of early development pathways in peacock suggest the adaptive divergence of the early development processes, including feather, bone and skeletomuscle development.

#### Peacock feathers: Clues from early development genes

Among the distinctive features of a peacock, the large and decorative feathers attract the most attention; particularly the long train, which is useful for their courtship behavior. The feather development in birds is primarily guided by the continuous reciprocal interactions between the epithelium and mesenchyme (Chuong et al., 2000). The analysis of the curated set of 2,146 feather-related genes (**Supplementary Note**) involved in feather development revealed that the activators of feather development including FGF, Wnt/β-catenin and TGF-β and, the inhibitors such as BMP and Notch-delta showed sequence divergence in peacock in comparison to the other bird genomes. The observed divergence in genes related to feather development provides useful genomic clues for the peculiar patterning and structure of peacock feathers.

### Adaptive Evolution in Immune-related Genes

In birds, the rate of sequence divergence in immune-related genes is usually higher than the other genes primarily due to the co-evolution of host-pathogen interactions (Ekblom et al., 2010). Several genes involved in the development of immune system and modulation of immune response have shown sequence divergence and signs of adaptive evolution in the peacock genome.

Multiple components of the innate immune system such as complement system and pathogen recognition system showed adaptive evolution. The C5 protein involved in the recruitment of cellular component of the immune system at the site of infection showed five unique substitutions. The α-subunit of C8 protein involved in forming the membrane attack complex (MAC)(Serna et al., 2016) showed MSA. Additionally, the CSF-1R gene, which is crucial for macrophage survival, differentiation, and proliferation (Pixley and Stanley, 2004), showed positive selection with positively selected sites and unique substitutions. Different components of NF-ĸB signaling such as MYD88, TRADD, SIGIRR, MAP3K14 and TLR5, which regulate the immune response against infections (Kaisho and Akira, 2006), showed signs of adaptations. The MYD88 protein, which is a part of Toll-like receptors (TLRs) mediated signaling, showed MSA and higher divergence from chicken in comparison to turkey among the species of the Galliformes order. Similarly, the genes TRADD, SIGIRR, MAP3K14, and TLR5 showed multiple unique substitutions. Furthermore, the pattern recognition receptors such as NLRC3, which regulates innate immune response by interacting with stimulators of interferon genes (Zhang et al., 2014), showed positive selection with positively selected sites and unique substitution.

Several genes regulating the T and B-cell response of the adaptive immune system also displayed adaptive evolution in peacock. The SPI-1 gene involved in B and T cell development by regulating the expression as well as alternative splicing of target genes (Hallier et al., 1998) showed MSA. The ITGAV and AQP3 genes, which are involved in T-cell movement and migration, showed unique substitutions and higher (2X) divergence from chicken as compared to turkey. Furthermore, different T-cell receptors and signaling proteins involved in T-cell activation such as SDC4, FLT4, NFATC3, and IL12B subunit showed sequence divergence and multiple signs of adaptation in peacock. CTLA4 gene, which is a negative regulator of T-cell response (Walunas et al., 1994), also showed multiple unique substitutions. A few other regulator genes of immune response also showed multiple signs of adaptation in peacock and are discussed in Supplementary Note. In addition, the gene family SSC4D involved in the development of immune system and the regulation of both innate and adaptive immunity (Asratian and Vasil’eva, 1976) showed expansion in peacock in comparison to chicken (**Supplementary Table 8**).

Taken together, it appears that the adaptive evolution of immune-related genes in peacock has occurred primarily in the components of innate immunity such as complement system, pattern recognition receptors, and monocyte development, and in the components of adaptive immunity such as T-cell response. It suggests that the immune system-related genes in peacock genome have significantly evolved to provide a selective advantage in fighting against infections.

### Body Dimensions

Follicle stimulating hormone receptor (FSHR), which is involved in regulating the cell growth, differentiation, and body dimensions of birds via cAMP-mediated PI3K-AKT and SRC-ERK1/2 signaling (Fayeye et al., 2006), showed multiple unique substitutions. Several genes such as MMP2, BMP7, TRAF6, TNF3, Neurochondrin, IGF, and NOX4, regulating bone morphogenesis and development in birds showed divergence as well as adaptive evolution in peacock. These genes primarily function as ligands or receptors for Wnt-beta-catenin, TGF-beta, p70S6K and PEDF signaling pathways. From these observations, it appears that the adaptive evolution of intracellular signaling and early development genes, which play significant roles in bone and skeletal muscle development, are perhaps beneficial for supporting its body dimensions.

### MSA genes involved in other cellular processes

Among the other genes that displayed multiple signs of adaptation, BRCA2, DNA-PKcs, FANCC, and INO80 genes were involved in the DNA double-strand break repair and recombination, FBXO15, USP53, and PSMD1-26S were part of ubiquitin-proteasomal protein degradation system, HERPUD1 and HSP90B1 genes were involved in stress response, and METTL5 gene had protein methyltransferase activity. Thus, DNA repair and protein turnover and modification were among the other cellular processes where a notable number of genes showed MSA.

## DISCUSSION

The most significant results emerged from the adaptive sequence divergence analysis, where a major fraction of genes involved in early development and immune system showed multiple signs of adaptive evolution (**Figure 4**). Similarly, the genes involved in the early development of feathers showed signs of adaptive evolution in the feather-specific gene set. In addition, the adaptive divergence observed in the genes involved in bone morphogenesis and skeletal muscle development perhaps explain the large body dimensions, stronger legs and spurs, and the ability to take short flights despite of a long train. Taken together, the evolution in the early development genes emerges as a prominent factor for explaining the molecular basis of the phenotypic evolution for Indian peacock.

**Figure 4:**
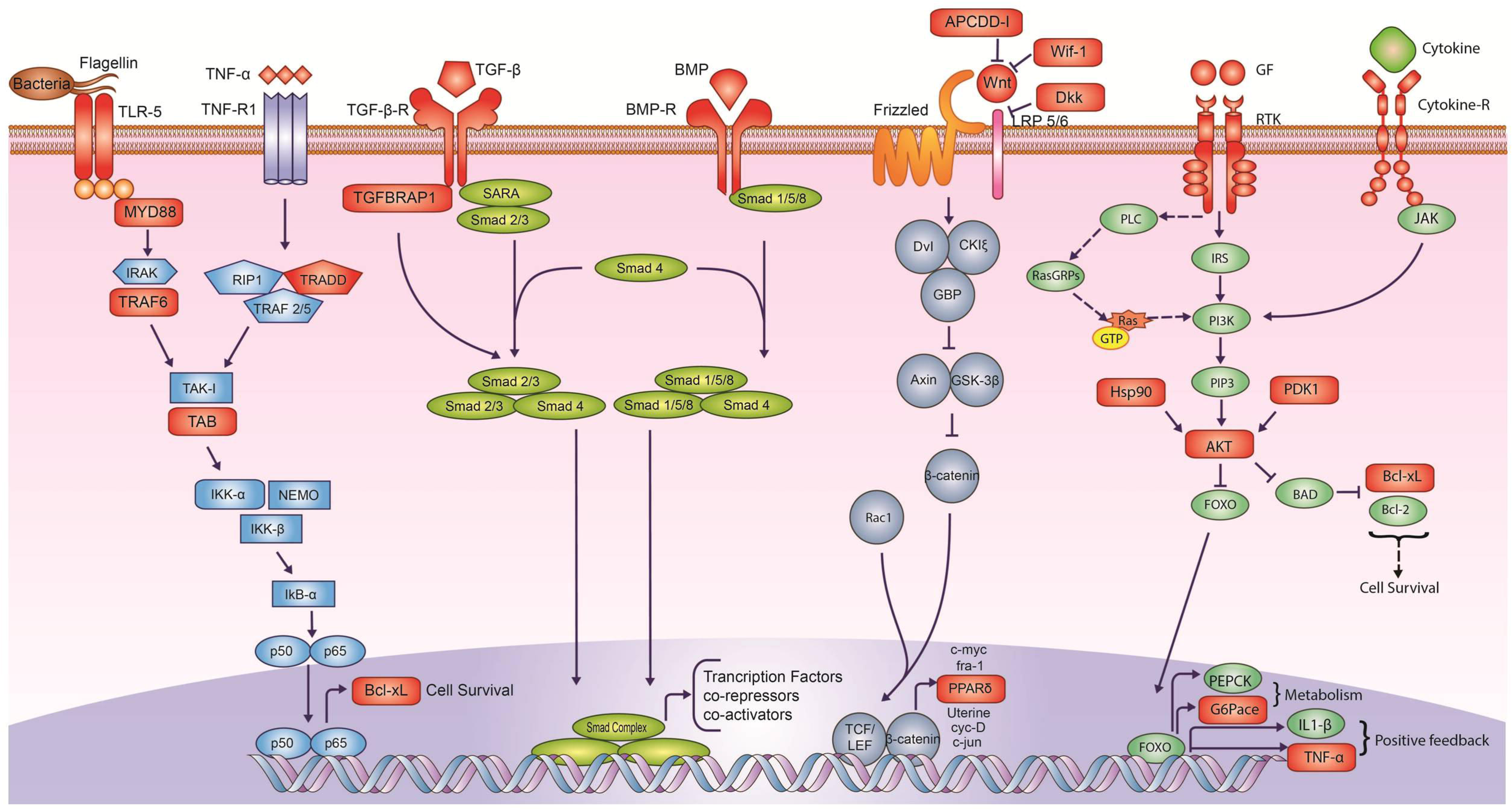
Adaptively evolved signaling pathways in peacock genome. The genes highlighted in Red colour showed signs of adaptive evolution such as positive selection and unique substitution. It is apparent that the receptors, ligands and regulators of early development pathways such as Wnt, TGF-β and BMP, showed adaptive sequence divergence in peacock. In the case of NF-KB, cytokine and growth factor signaling pathways, the proteins involved in intermediate signal transduction also showed adaptive sequence divergence. Individual pathways are colour coded separately.

Though birds are the natural host of viruses and are also prone to avian viral infections (Alexander, 2000;Berg, 2000;Liu et al., 2005), peacocks have a longer average life span, and are also found to be resistant to the new viral strain pathogenic to chicken and turkey (Sun et al., 2007), pointing towards the presence of a robust immune system. The strong immunity against pathogens and infections could be attributed to the adaptive divergence observed in the components of the innate immune system (complement and pathogen recognition system), adaptive immune response (B and T cell development), and other genes responsible for the overall immune system development. The adaptive evolution observed for immune genes in peacock appears to be indicative of a higher parasite load consistent with Hamilton-Zuk hypothesis (Balenger and Zuk, 2014). Though the results were obtained from the comparative genomic analysis of peacock, some of the insights may also be applicable to the other related species in the pheasant group. The comparative genomic analysis presented in this work provides novel insights on the phenotypic evolution of Indian Peacock and the genomic clues from this study will serve as leads for further studies to decipher the genotype-phenotype interactions for peacock. In addition, this study will also help in devising better strategies for the management and conservation of peacock population, which is showing a decline mainly because of habitat deterioration, poaching for train-feathers, use of pesticides and chemical fertilizers.

## Acknowledgements

We thank Dr. Atul Gupta, Wildlife Veterinary Officer, Van Vihar National Park, Bhopal and Director, Van Vihar National Park, Bhopal, India for providing the blood samples of peacock. We also acknowledge the help of Dr. Tista Joseph and Dr. Niraj Dahe, Wildlife Veterinary Officers (Wildlife SOS India) at Van Vihar National Park for carrying out the sample collection procedure. We thank the HPC facility and NGS facility at IISER Bhopal. The authors SKJ, AG and RS thank the Department of Science and Technology for the DST-INSPIRE fellowship. We also thank the intramural research funds provided by IISER Bhopal.

## Contributions

VKS conceived and coordinated the project. RS prepared the DNA samples, performed sequencing and the molecular sexing assay. AG performed the *de novo* and reference-based genome assembly. PM, AKS, AG and SKJ performed the genome annotations. SKJ and PM performed the phylogenetic tree analyses. SKJ performed the dN/dS, positive selection, and statistical analysis. SKJ, AG and AR performed the unique substitution and SIFT analyses. PM performed the gene gain/loss analysis. SKJ, VPPK and AG created figures. SKJ, AG, VKS, NV, and AS analysed the data and wrote the manuscript. All the authors have read and approved the final manuscript.

## REFERENCES

Alexander, D.J. (2000). A review of avian influenza in different bird species. Vet Microbiol 74, 3–13.

Asratian, A.A., and Vasil’;eva, V.I. (1976). [Immunological structure of the population of Erevan with regard to Mycoplasma hominis]. Zh Eksp Klin Med 16, 59–61.

Balenger, S.L., and Zuk, M. (2014). "Testing the Hamilton–Zuk hypothesis: past, present, and future". The Society for Integrative and Comparative Biology).

Berg, T.P. (2000). Acute infectious bursal disease in poultry: a review. Avian Pathol 29, 175–194.

Bintanja, R., and Van De Wal, R. (2008). North American ice-sheet dynamics and the onset of 100,000-year glacial cycles. Nature 454, 869–872.

Bornelöv, S., Seroussi, E., Yosefi, S., Pendavis, K., Burgess, S.C., Grabherr, M., Friedman-Einat, M., and Andersson, L. (2017). Correspondence on Lovell et al.: identification of chicken genes previously assumed to be evolutionarily lost. Genome biology 18, 112.

Brack, A.S., Conboy, M.J., Roy, S., Lee, M., Kuo, C.J., Keller, C., and Rando, T.A. (2007). Increased Wnt signaling during aging alters muscle stem cell fate and increases fibrosis. Science 317, 807–810.

Chuong, C.M., Chodankar, R., Widelitz, R.B., and Jiang, T.X. (2000). Evo-devo of feathers and scales: building complex epithelial appendages. Curr Opin Genet Dev 10, 449–456.

Crowe, R., Henrique, D., Ish-Horowicz, D., and Niswander, L. (1998). A new role for Notch and Delta in cell fate decisions: patterning the feather array. Development 125, 767–775.

Denton, J.F., Lugo-Martinez, J., Tucker, A.E., Schrider, D.R., Warren, W.C., and Hahn, M.W. (2014). Extensive error in the number of genes inferred from draft genome assemblies. PLoS computational biology 10, e1003998.

Edgar, R.C. (2004). MUSCLE: multiple sequence alignment with high accuracy and high throughput. Nucleic Acids Res 32, 1792–1797.

Ekblom, R., French, L., Slate, J., and Burke, T. (2010). Evolutionary analysis and expression profiling of zebra finch immune genes. Genome Biol Evol 2, 781–790.

Fayeye, T., Ayorinde, K., Ojo, V., and Adesina, O. (2006). Frequency and influence of some major genes on body weight and body size parameters of Nigerian local chickens. Livestock research for rural development 18, 37.

Groppe, J., Greenwald, J., Wiater, E., Rodriguez-Leon, J., Economides, A.N., Kwiatkowski, W., Affolter, M., Vale, W.W., Izpisua Belmonte, J.C., and Choe, S. (2002). Structural basis of BMP signalling inhibition by the cystine knot protein Noggin. Nature 420, 636–642.

Guindon, S., Dufayard, J.F., Lefort, V., Anisimova, M., Hordijk, W., and Gascuel, O. (2010). New algorithms and methods to estimate maximum-likelihood phylogenies: assessing the performance of PhyML 3.0. Syst Biol 59, 307–321.

Hallier, M., Lerga, A., Barnache, S., Tavitian, A., and Moreau-Gachelin, F. (1998). The transcription factor Spi-1/PU.1 interacts with the potential splicing factor TLS. J Biol Chem 273, 4838–4842.

Han, M.V., Thomas, G.W., Lugo-Martinez, J., and Hahn, M.W. (2013). Estimating gene gain and loss rates in the presence of error in genome assembly and annotation using CAFE 3. Mol Biol Evol 30, 1987–1997.

Hedges, S.B., Dudley, J., and Kumar, S. (2006). TimeTree: a public knowledge-base of divergence times among organisms. Bioinformatics 22, 2971–2972.

Huang, Y., Li, Y., Burt, D.W., Chen, H., Zhang, Y., Qian, W., Kim, H., Gan, S., Zhao, Y., and Li, J. (2013). The duck genome and transcriptome provide insight into an avian influenza virus reservoir species. Nature genetics 45, 776.

Huerta-Cepas, J., Szklarczyk, D., Forslund, K., Cook, H., Heller, D., Walter, M.C., Rattei, T., Mende, D.R., Sunagawa, S., Kuhn, M., Jensen, L.J., Von Mering, C., and Bork, P. (2016). eggNOG 4.5: a hierarchical orthology framework with improved functional annotations for eukaryotic, prokaryotic and viral sequences. Nucleic Acids Res 44, D286–293.

Huxley, T.H. (1968). On the origin of species. University of Michigan P.

Kaiser, V.B., Van Tuinen, M., and Ellegren, H. (2007). Insertion events of CR1 retrotransposable elements elucidate the phylogenetic branching order in galliform birds. Mol Biol Evol 24, 338–347.

Kaisho, T., and Akira, S. (2006). Toll-like receptor function and signaling. J Allergy Clin Immunol 117, 979–987; quiz 988.

Kan, X.-Z., Li, X.-F., Lei, Z.-P., Chen, L., Gao, H., Yang, Z.-Y., Yang, J.-K., Guo, Z.-C., Yu, L., and Zhang, L.-Q. (2010). Estimation of divergence times for major lineages of galliform birds: evidence from complete mitochondrial genome sequences. African Journal of Biotechnology 9, 3073–3078.

Kanehisa, M., and Goto, S. (2000). KEGG: kyoto encyclopedia of genes and genomes. Nucleic Acids Res 28, 27–30.

Kimball, R.T., Braun, E.L., Zwartjes, P.W., Crowe, T.M., and Ligon, J.D. (1999). A molecular phylogeny of the pheasants and partridges suggests that these lineages are not monophyletic. Mol Phylogenet Evol 11, 38–54.

Klaus, A., and Birchmeier, W. (2008). Wnt signalling and its impact on development and cancer. Nat Rev Cancer 8, 387–398.

Kumar, P., Henikoff, S., and Ng, P.C. (2009). Predicting the effects of coding non-synonymous variants on protein function using the SIFT algorithm. Nat Protoc 4, 1073–1081.

Li, H., and Durbin, R. (2011). Inference of human population history from individual whole-genome sequences. Nature 475, 493–496.

Lim, X., and Nusse, R. (2013). Wnt signaling in skin development, homeostasis, and disease. Cold Spring Harb Perspect Biol 5.

Liu, J., Xiao, H., Lei, F., Zhu, Q., Qin, K., Zhang, X.W., Zhang, X.L., Zhao, D., Wang, G., Feng, Y., Ma, J., Liu, W., Wang, J., and Gao, G.F. (2005). Highly pathogenic H5N1 influenza virus infection in migratory birds. Science 309, 1206.

Lovell, P.V., Wirthlin, M., Wilhelm, L., Minx, P., Lazar, N.H., Carbone, L., Warren, W.C., and Mello, C.V. (2014). Conserved syntenic clusters of protein coding genes are missing in birds. Genome biology 15, 565.

Loveridge, N., Farquharson, C., Hesketh, J.E., Jakowlew, S.B., Whitehead, C.C., and Thorp, B.H. (1993). The control of chondrocyte differentiation during endochondral bone growth in vivo: changes in TGF-beta and the proto-oncogene c-myc. J Cell Sci 105 ( Pt 4), 949–956.

Loyau, A., Jalme, M.S., and Sorci, G. (2005a). Intra- and Intersexual Selection for Multiple Traits in the Peacock (Pavo cristatus). Ethology 111, 810–820.

Loyau, A., Saint Jalme, M., Cagniant, C., and Sorci, G. (2005b). Multiple sexual advertisements honestly reflect health status in peacocks (Pavo cristatus). Behavioral Ecology and Sociobiology 58, 552–557.

Moyle, W.R., Campbell, R.K., Myers, R.V., Bernard, M.P., Han, Y., and Wang, X. (1994). Co-evolution of ligand-receptor pairs. Nature 368, 251–255.

Nadachowska-Brzyska, K., Li, C., Smeds, L., Zhang, G., and Ellegren, H. (2015). Temporal dynamics of avian populations during Pleistocene revealed by whole-genome sequences. Current Biology 25, 1375–1380.

Nadachowska-Brzyska, K., Burri, R., Smeds, L., and Ellegren, H. (2016). PSMC analysis of effective population sizes in molecular ecology and its application to black-and-white Ficedula flycatchers. Molecular Ecology 25, 1058–1072.

Nie, X., Luukko, K., and Kettunen, P. (2006). BMP signalling in craniofacial development. Int J Dev Biol 50, 511–521.

Nishimura, R., Hata, K., Matsubara, T., Wakabayashi, M., and Yoneda, T. (2012). Regulation of bone and cartilage development by network between BMP signalling and transcription factors. J Biochem 151, 247–254.

O’ leary, N.A., Wright, M.W., Brister, J.R., Ciufo, S., Haddad, D., Mcveigh, R., Rajput, B., Robbertse, B., Smith-White, B., Ako-Adjei, D., Astashyn, A., Badretdin, A., Bao, Y., Blinkova, O., Brover, V., Chetvernin, V., Choi, J., Cox, E., Ermolaeva, O., Farrell, C.M., Goldfarb, T., Gupta, T., Haft, D., Hatcher, E., Hlavina, W., Joardar, V.S., Kodali, V.K., Li, W., Maglott, D., Masterson, P., Mcgarvey, K.M., Murphy, M.R., O’neill, K., Pujar, S., Rangwala, S.H., Rausch, D., Riddick, L.D., Schoch, C., Shkeda, A., Storz, S.S., Sun, H., Thibaud-Nissen, F., Tolstoy, I., Tully, R.E., Vatsan, A.R., Wallin, C., Webb, D., Wu, W., Landrum, M.J., Kimchi, A., Tatusova, T., Dicuccio, M., Kitts, P., Murphy, T.D., and Pruitt, K.D. (2016). Reference sequence (RefSeq) database at NCBI: current status, taxonomic expansion, and functional annotation. Nucleic Acids Res 44, D733–745.

Ouyang, Y.-N., Yang, Z.-Y., Li, D.-L., Huo, J.-L., Qian, K., and Miao, Y.-W. (2009). Genetic Divergence between Pavo muticus and Pavo cristatus by Cyt b Gene. Journal of Yunnan Agricultural University 2, 014.

Owens, I.P., and Short, R.V. (1995). Hormonal basis of sexual dimorphism in birds: implications for new theories of sexual selection. Trends Ecol Evol 10, 44–47.

Pixley, F.J., and Stanley, E.R. (2004). CSF-1 regulation of the wandering macrophage: complexity in action. Trends Cell Biol 14, 628–638.

Pownall, M.E., and Isaacs, H.V. (Year). “Fgf signalling in vertebrate development”, in: Colloquium Series on Developmental Biology: Morgan & Claypool Life Sciences), 1–75.

Ramesh, K., and Mcgowan, P. (2009). On the current status of Indian peafowl Pavo cristatus (Aves: Galliformes: Phasianidae): keeping the common species common. Journal of Threatened Taxa 1, 106–108.

Rice, P., Longden, I., and Bleasby, A. (2000). EMBOSS: the European Molecular Biology Open Software Suite. Trends Genet 16, 276–277.

Serna, M., Giles, J.L., Morgan, B.P., and Bubeck, D. (2016). Structural basis of complement membrane attack complex formation. Nat Commun 7, 10587.

Stock, A.D., and Bunch, T.D. (1982). The evolutionary implications of chromosome banding pattern homologies in the bird order Galliformes. Cytogenet Cell Genet 34, 136–148.

Sun, L., Zhang, G.H., Jiang, J.W., Fu, J.D., Ren, T., Cao, W.S., Xin, C.A., Liao, M., and Liu, W.J. (2007). A Massachusetts prototype like coronavirus isolated from wild peafowls is pathogenic to chickens. Virus Res 130 121–128.

Vilella, A.J., Severin, J., Ureta-Vidal, A., Heng, L., Durbin, R., and Birney, E. (2009). EnsemblCompara GeneTrees: Complete, duplication-aware phylogenetic trees in vertebrates. Genome Res 19, 327–335.

Walunas, T.L., Lenschow, D.J., Bakker, C.Y., Linsley, P.S., Freeman, G.J., Green, J.M., Thompson, C.B., and Bluestone, J.A. (1994). CTLA-4 can function as a negative regulator of T cell activation. Immunity 1, 405–413.

Wang, N., Kimball, R.T., Braun, E.L., Liang, B., and Zhang, Z. (2013). Assessing phylogenetic relationships among galliformes: a multigene phylogeny with expanded taxon sampling in Phasianidae. PLoS One 8, e64312.

Wright, A.E., Harrison, P.W., Zimmer, F., Montgomery, S.H., Pointer, M.A., and Mank, J.E. (2015). Variation in promiscuity and sexual selection drives avian rate of Faster-Z evolution. Molecular Ecology 24, 1218–1235.

Yang, Z. (2007). PAML 4: phylogenetic analysis by maximum likelihood. Mol Biol Evol 24, 1586–1591.

Yang, Z., Wong, W.S., and Nielsen, R. (2005). Bayes empirical bayes inference of amino acid sites under positive selection. Mol Biol Evol 22, 1107–1118.

Zahavi, A. (1975). Mate selection—A selection for a handicap. Journal of Theoretical Biology 53, 205–214.

Zhang, J., Nielsen, R., and Yang, Z. (2005). Evaluation of an improved branch-site likelihood method for detecting positive selection at the molecular level. Mol Biol Evol 22, 2472–2479.

Zhang, L., Mo, J., Swanson, K.V., Wen, H., Petrucelli, A., Gregory, S.M., Zhang, Z., Schneider, M., Jiang, Y., and Fitzgerald, K.A. (2014). NLRC3, a member of the NLR family of proteins, is a negative regulator of innate immune signaling induced by the DNA sensor STING. Immunity 40, 329–341.

